# Locomotor suppression by a monosynaptic amygdala to brainstem circuit

**DOI:** 10.1101/724252

**Authors:** Thomas K. Roseberry, Arnaud L. Lalive, Benjamin D. Margolin, Anatol C. Kreitzer

## Abstract

The control of locomotion is fundamental to vertebrate animal survival. Defensive situations require an animal to rapidly decide whether to run away or suppress locomotor activity to avoid detection. While much of the neural circuitry involved in defensive action selection has been elucidated, top-down modulation of brainstem locomotor circuitry remains unclear. Here we provide evidence for the existence and functionality of a monosynaptic connection from the central amygdala (CeA) to the mesencephalic locomotor region (MLR) that inhibits locomotion in unconditioned and conditioned defensive behavior in mice. We show that locomotion stimulated by airpuff coincides with increased activity of MLR glutamatergic neurons. Using retrograde tracing and *ex vivo* electrophysiology, we find that the CeA makes a monosynaptic connection with the MLR. In the open field, *in vivo* stimulation of this projection suppressed spontaneous locomotion, whereas inhibition of this projection had no effect. However, inhibiting CeA terminals within the MLR increased both neural activity and locomotor responses to airpuff. Finally, using a conditioned avoidance paradigm known to activate CeA neurons, we find that inhibition of the CeA projection increased successful escape, whereas activating the projection reduced escape. Together these results provide evidence for a new circuit substrate influencing locomotion and defensive behaviors.

## Introduction

Locomotion is a fundamental aspect of defensive behavior. When an animal perceives a threat, competitive neural processes determine whether to freeze for detection avoidance or to escape (Fadok et al., 2017; Herry and Johansen, 2014; LeDoux, 2000). Locomotor circuitry must rapidly and efficiently respond if the animal is to escape. Conversely, if the animal is to freeze then this potent locomotor response must be aborted. Thus, the control of locomotion is a crucial aspect of the fear response.

The mesencephalic locomotor region (MLR) MLR is an evolutionarily-conserved brainstem region that links higher order brain regions in the central nervous system to central pattern generators in the spinal cord that govern locomotion (Grillner et al., 2008; Ryczko and Dubuc, 2013; Sherman et al., 2015). The MLR was originally defined in cats, where electrical stimulation can drive running at short latencies (Shik et al., 1966). More recently, glutamatergic neurons in the mouse MLR were shown to be necessary and sufficient for locomotion (Lee et al., 2014; Roseberry et al., 2016). Given the key role of the MLR in locomotion, it is likely that defensive locomotion is also coordinated via the MLR, although the precise circuit substrates are still being worked out (Hormigo et al., 2019).

The amygdaloid complex plays a major role in defensive behaviors (Ciocchi et al., 2010; Herry and Johansen, 2014; LeDoux, 2000; Pare et al., 2004; Tovote et al., 2016). The main output nucleus of the amygdaloid complex is the CeA which modulates autonomic attributes of the defensive response, including heart and breathing rates (Kapp et al., 1979; LeDoux et al., 1988), in addition to impacting the decision whether to escape or freeze (Fadok et al., 2017; Tovote et al., 2016). Lesion of the CeA decreases freezing (Ciocchi et al., 2010; LeDoux et al., 1988) and increases escape responses to learned (Choi and Brown, 2003; Choi et al., 2010) and innately dangerous stimuli (Choi and Kim, 2010), but lesions of the other amygdaloid nuclei individually do not have the same effect (Choi et al., 2010). This suggests that, among the amygdaloid nuclei, the CeA is especially important for shaping freezing-vs-escape responses. Recent work has begun to elucidate the cell types within the CeA by demonstrating that activation of somatostatin neurons can stop locomotion and avoidance responses (Fadok et al., 2017; Yu et al., 2016). These results suggest that the CeA can inhibit locomotion in defensive situations.

We recently reported that neurons within the CeA project to glutamatergic neurons in the MLR (Roseberry et al., 2016). Although previously observed (Ciocchi et al., 2010; Hopkins and Holstege, 1978; Liang et al., 2015), this potentially critical projection has been largely overlooked because it was thought that the labeling in the CeA arose mainly from fibers that terminate in the periaqueductal gray (PAG) and pontine nuclei (Ciocchi et al., 2010; Oka et al., 2008). Here, we test the hypothesis that this connection could play an important role in suppression of locomotion during periods of fear.

## Results

### Functionality of the CeA-MLR projection

To determine how locomotion-encoding neurons within the MLR respond to a noxious stimulus, we performed acute, head-fixed, single-unit recordings from the MLR of awake mice while monitoring locomotion on a circular treadmill (Figure 1A-C). Delivery of brief air puffs to the face triggered brief bouts of running (Figure 1A, bottom) that were associated with a transient increase in MLR neuron activity (Figure 1C,D). Because glutamatergic neurons are the main cell type driving locomotion in the MLR (Roseberry et al., 2016), we repeated these experiments in an additional set of vGluT2-cre mice that had been previously injected with DIO-ChR2 to permit optogenetic identification of glutamatergic neurons (Roseberry et al., 2016). Remarkably, 100% (14 of 14) of optogenetically identified glutamatergic neurons in the MLR neurons significantly increased firing in response to airpuff (Figure 1E-H and Supplemental Figure 1). Furthermore, elevated activity of MLR glutamatergic neurons prior to the onset of the airpuff in stationary animals was associated with an increased probability that those animals would start to run within 1 second following airpuff onset (Figure 1I-J).

**Figure 1.**
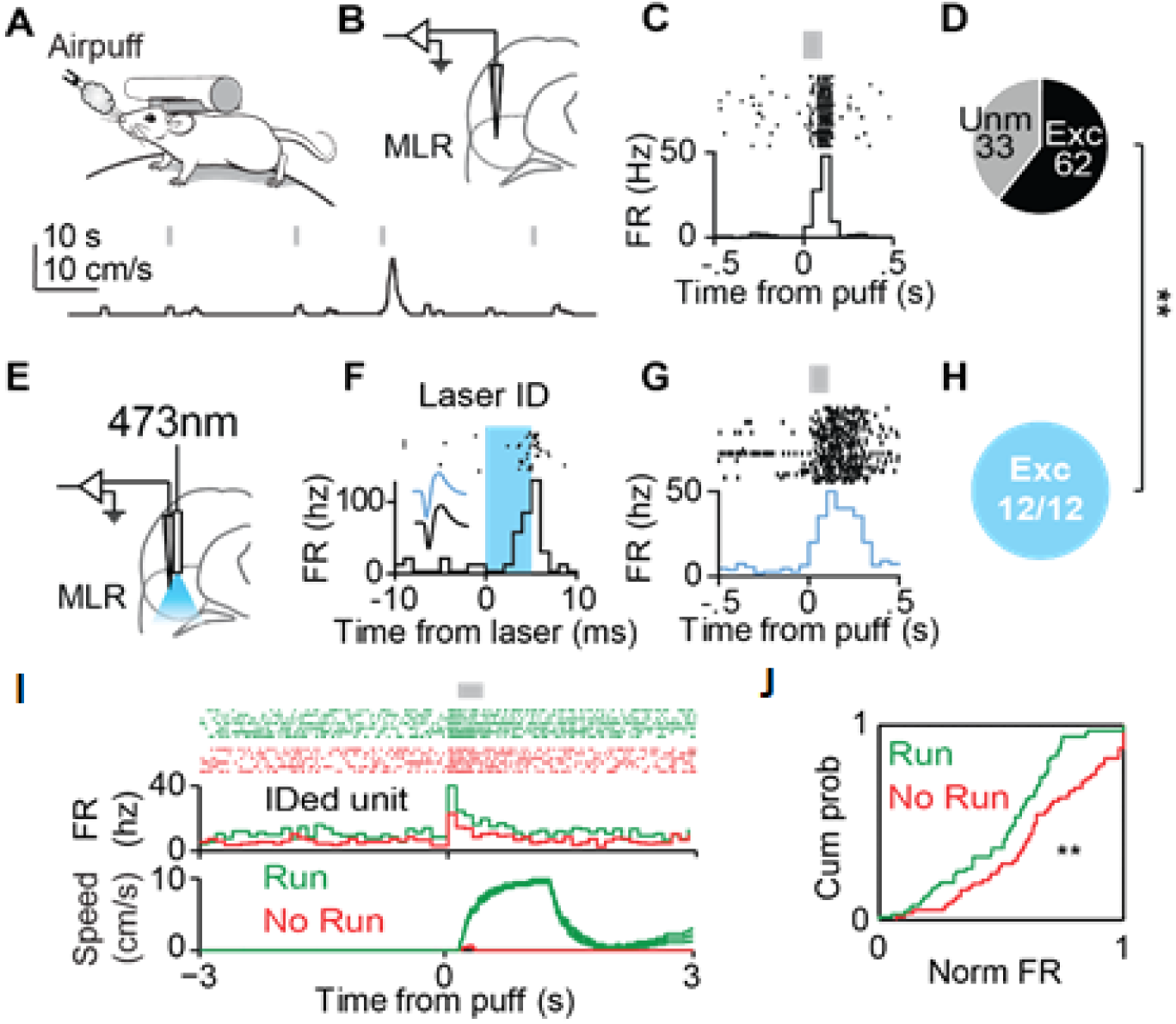
Aversive airpuff drives locomotion and MLR glutamatergic neuron activity. (A) Schematic for recording MLR neurons and locomotion during airpuff (top). Exemple locomotor trace from circular treadmill (bottom). Gray lines indicate airpuff delivery. (B) Targeting of MLR for single unit recordings. (C) Raster (top) and PSTH (bottom) of example MLR neuron excited by airpuff. (D) Proportion of significantly excited or unmodulated MLR neurons during airpuff (n = 95 neurons from 4 mice). (E) Schematic for recording optogenetically identified MLR glutamatergic neurons. (F) Raster and PSTH of an example identified MLR glutamatergic neuron during laser stimulation (shaded blue region). Inset, spike waveforms during (blue) and outside (black) of laser pulse window. (G) Raster and PSTH of neuron in Panel H in response to airpuff (gray box). (H) Fraction of identified neurons excited by airpuff (** p < .01, χ2 test, n = 12 neurons from 5 mice). (I) Top and middle, raster and PSTH, respectively of identified unit during trials in which the mouse ran (green) and trials in which the mouse did not run (red). Bottom, average speed traces for both response types. (J) Cumulative distribution plots for running (green) and no running (red) trials (p < 0.01, Kolmogrov Smirnov test with matched sample sizes). Firing rate normalized to the maximum firing during airpuff.

We hypothesized that input from the CeA could be ideally positioned to modulate MLR activity and potentially locomotor responses to aversive stimuli. Therefore, we first sought to confirm the CeA-MLR projection by injecting retrograde CAV2-Cre virus (Junyent and Kremer, 2015) in the MLR and AAV5-DIO-ChR2-eYFP in the CeA (Figure 2A). Cell bodies were labeled with eYFP in both the centrolateral amygdala (CeL) and centromedial amygdala (CeM), the two divisions of the CeA (Figure 1B), and eYFP-positive fibers were observed across the MLR (Figure 2B).

**Figure 2.**
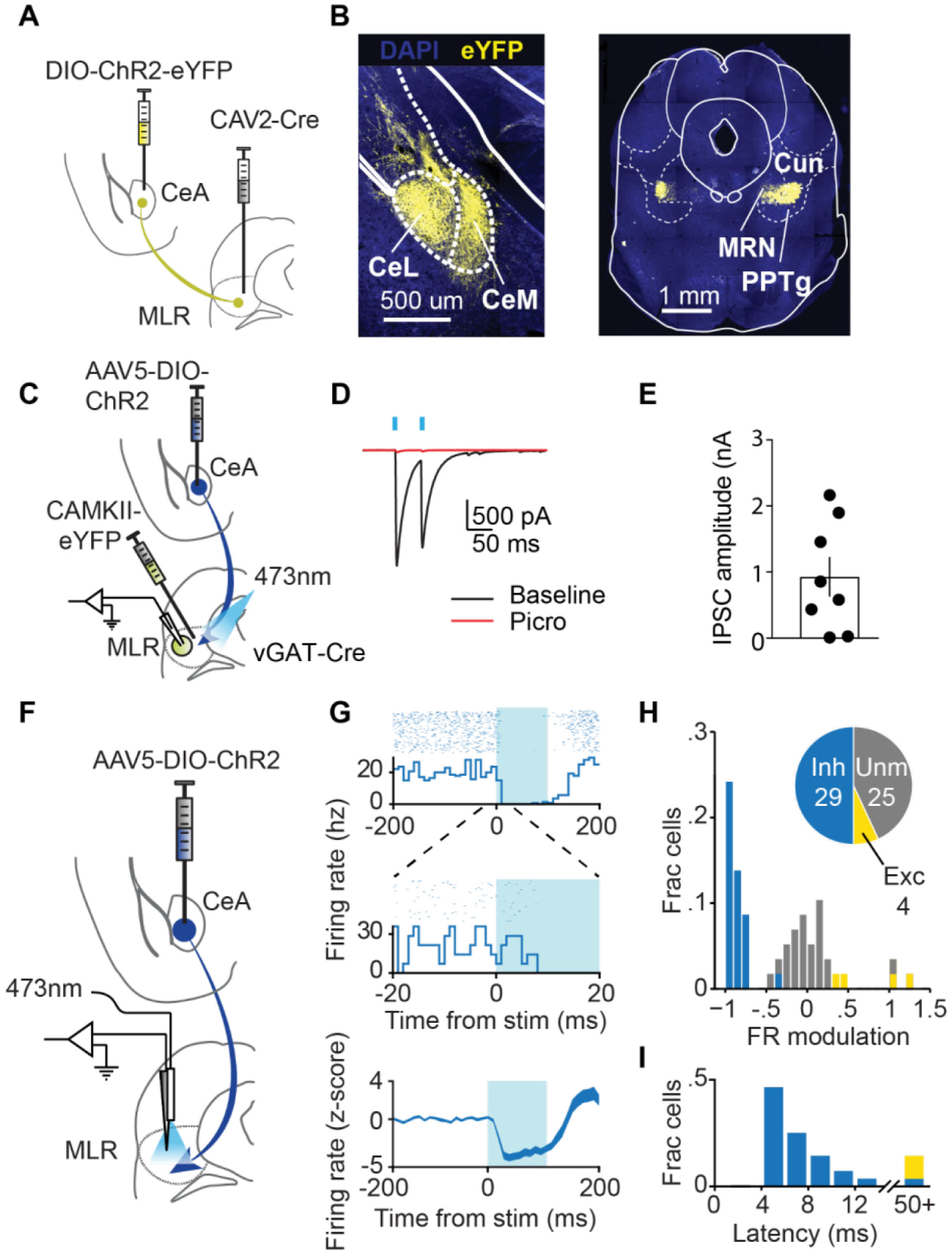
The CeA makes GABAergic synapses with MLR neurons. (A) Schematic for labeling MLR-projecting CeA cells and fibers. (B) Left, example of cells labelled in the centrolateral amygdala (CeL) and centromedial amygdala (CeM). Right, example of CeA fibers in the MLR. Cun, cuneiform nucleus; MRN, mesencephalic reticular nucleus; PPTg, pedunculopontine tegmental nucleus. (C) Schematic for exciting CeA terminals in the MLR in slice while patching identified glutamatergic cells. (D) Example traces of IPSCs in MLR glutamatergic neurons during optical stim (blue boxes) before (black) and after (red) picrotoxin infusion. (E) Summary of IPSC amplitudes during CeA terminal stimulation. (F) Schematic of strategy for exciting CeA terminals in the MLR while recording activity. (G) Top, example PSTH and raster of an inhibited MLR neuron. Middle, same neuron at smaller timescale. Bottom, time course of the population of significantly inhibited neurons. (H) Histogram of firing rate modulation. Inset, proportions of inhibited (blue, Inh), excited (yellow, Exc) and unmodulated (grey, Unm) neurons. (I) Histogram of latencies to significant modulation.

To test the functionality of the CeA-MLR projection, we performed *ex vivo* whole-cell patch-clamp recordings from identified glutamatergic cells in the MLR while stimulating CeA terminals (Figure 2C-E). To restrict viral expression to the GABAergic CeA (Oka et al., 2008) (Figure S1A) and prevent expression by projection neurons in neighboring regions, we injected a DIO-ChR2 into the CeA of vGAT-Cre mice and waited 4 to 6 weeks for viral expression and construct trafficking to synaptic terminals. CaMKII-eYFP was injected in the MLR to label glutamatergic cells (Lee et al., 2014; Roseberry et al., 2016). Blue light elicited IPSCs in 13 of 14 identified MLR glutamatergic cells (Figure 2D-E), and IPSCs were completely blocked with picrotoxin in 3/3 of cells (Figure 2D, F), indicating that the CeA projection is indeed GABAergic (Oka et al., 2008; Tovote et al., 2016).

We next examined how activation of CeA terminals modulates firing rates of MLR neurons *in vivo*.,After injecting DIO-ChR2 in the CeA of vGAT-Cre mice, we performed acute head-fixed recordings in the MLR using a silicon optrode probe (Figure 2F and Supplemental Figure 2A-C). Blue light was pulsed for 100 ms at 0.5 Hz while single units were recorded. Half of MLR neurons in these recordings were significantly inhibited at short latencies (mean latency 6.37 ± 0.43 ms, Figure 2F-H), confirming that monosynaptic inhibition from the CeA is sufficient to suppress MLR neuronal activity *in vivo*. In contrast, relatively few neurons were excited and the latency to significant excitation was much longer than the inhibitory responses, suggesting these excitatory responses are likely a result of polysynaptic disinhibition (Figure 2H and Supplemental Figure 2C).

We next tested whether the CeA-driven suppression of MLR neurons was sufficient to affect locomotion. As with the physiological recording experiments (Figure 2), DIO-ChR2 virus was injected into the CeA of vGAT-Cre mice (Figure 3A,B and Supplemental Figure 2D-E). During the same surgery, fiber optics were implanted bilaterally over the MLR and a headbar was affixed to the skull. After habituation to head-fixation, 40 Hz blue light stimulation (15 ms ON, 10ms OFF) was delivered in 5 second epochs to activate CeA terminals in the MLR. Stimulation onset was restricted to periods when the mouse was running at speeds > 5 cm/s to specifically test whether activation of the CeA projection to the MLR was sufficient to suppress ongoing locomotion. Strikingly, all mice immediately ceased running on all trials (Figure 3C-E). Anecdotally the mice did not freeze but instead appeared to elongate and flex thoracic muscles in an attempt to continue running. Although control mice did spontaneously stop locomoting during the light delivery period on a subset of trials (Figure 3E), these stops occurred at a longer latency than the optogenetically driven changes in locomotion (Figure 3G) and mice were significantly less likely to restart running during stimulation (Figure 3F) These results suggest that the CeA activation of sufficient to potently inhibit locomotion through a direct monosynaptic connection to the MLR.

**Figure 3.**
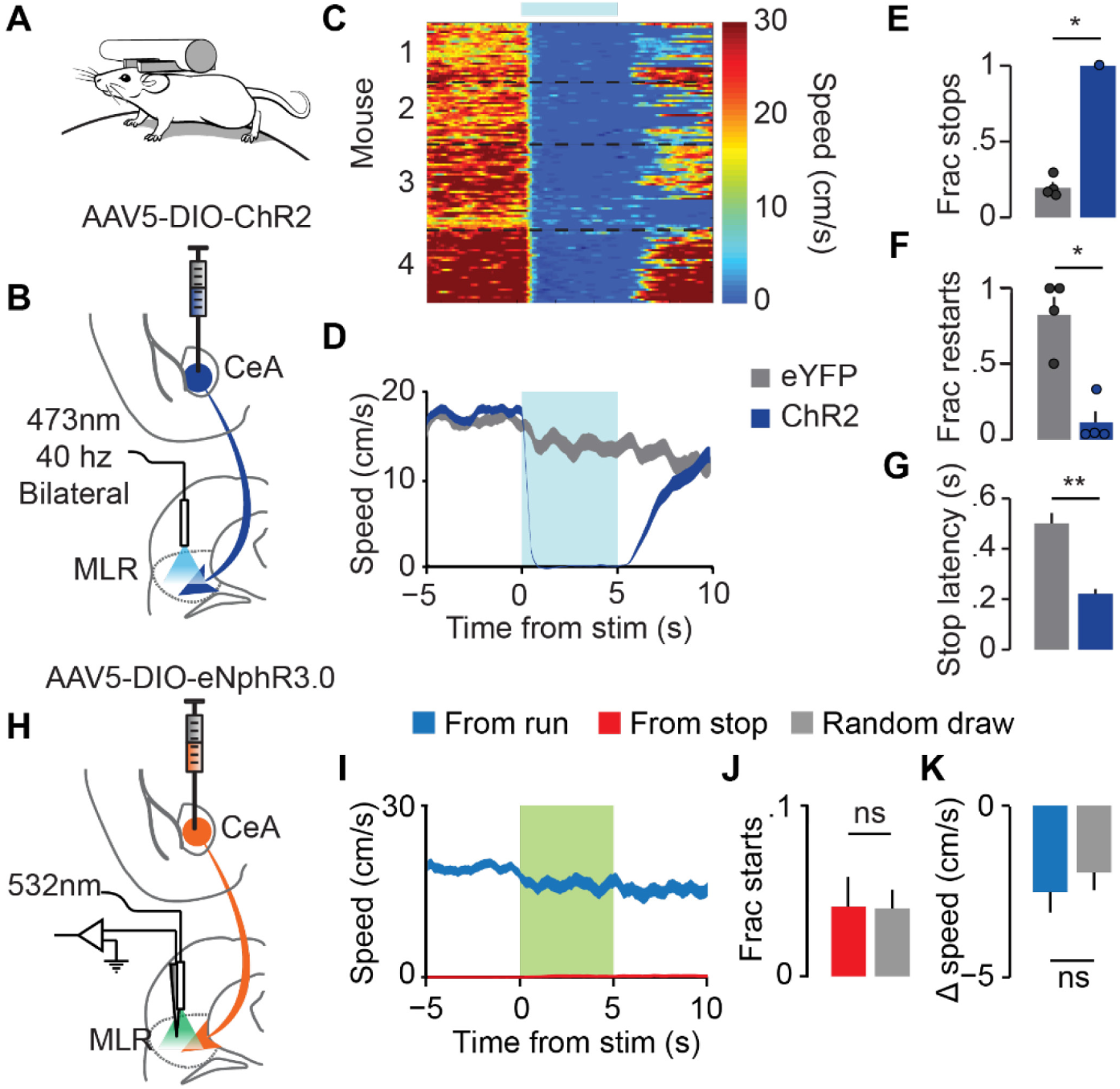
Activation of CeA terminals in MLR suppresses locomotion but silencing does not affect ongoing spontaneous movement. (A) Behavioral recording configuration and (B) schematic of strategy for exciting CeA terminals in the MLR during locomotor behavior. (C) Heat plot of mouse speed for all stimulation trials. Individual mice are separated by black dotted lines. (D) Population average for all trials in ChR2 mice (blue) and controls (grey). (E) Fraction of trials resulting in a stop. (F) Fraction of trials in which the mouse restarted after stop. (G) Latency to stop (*p < .05, **p < .01, rank-sum). (H) Schematic of strategy for inhibiting CeA terminals in the MLR. (I) Speed traces (left), fraction of starts made (middle), and average change in speed (right) during laser (green box) while the mouse was running (blue) or stopped (red). (J) Fraction of initially stationary trials in which the mouse initiated locomotion. Boxes with grey fill are from a random selection outside laser times with the same starting conditions. (K) Average change in speed in trials in which the mouse was initially running. (ns: p > 0.05, rank-sum).

We next tested if the CeA was tonically inhibiting locomotion through its connection with MLR neurons. vGAT-Cre mice were injected with DIO-eNpHR3.0 in the CeA (Figure 3H). After waiting 6 weeks for expression and transport to axon terminals, we recorded MLR neuronal responses during optogenetic inhibition of the GABAergic terminals from the CeA. Unlike with terminal excitation, we observed no change in speed or number of starts during optogenetic suppression of amygdala inputs, whether the mice were running or stationary (Figure 3H-K).

### The CeA-MLR connection inhibits locomotor responses to aversive stimuli

CeA projection neurons are active during airpuff (Cui et al., 2017). Therefore, we hypothesized that input from the CeA to the MLR could be responsible for differences in neural activity and behavior observed between running and not running in response to airpuff (Figure 1). To test this, we injected DIO-eNpHR3.0 into the CeA of vGAT-Cre mice and recorded activity of MLR neurons using acute, awake, head-fixed electrophysiological recordings during airpuff delivery (Figure 4 A-B). We then suppressed CeA projections to the MLR during airpuffs on alternating trials. In mice that were injected with DIO-eNpHR3.0 in the CeA, light delivery in the MLR resulted in a 31% increase in MLR firing during airpuff, relative to no laser trials (Figure 4C-D). In contrast, mice injected with eYFP control virus did not show a significant change in firing during laser trials (Figure 4D). These results are consistent with elevated activity in the CeA during an airpuff inhibiting the MLR response.

**Figure 4.**
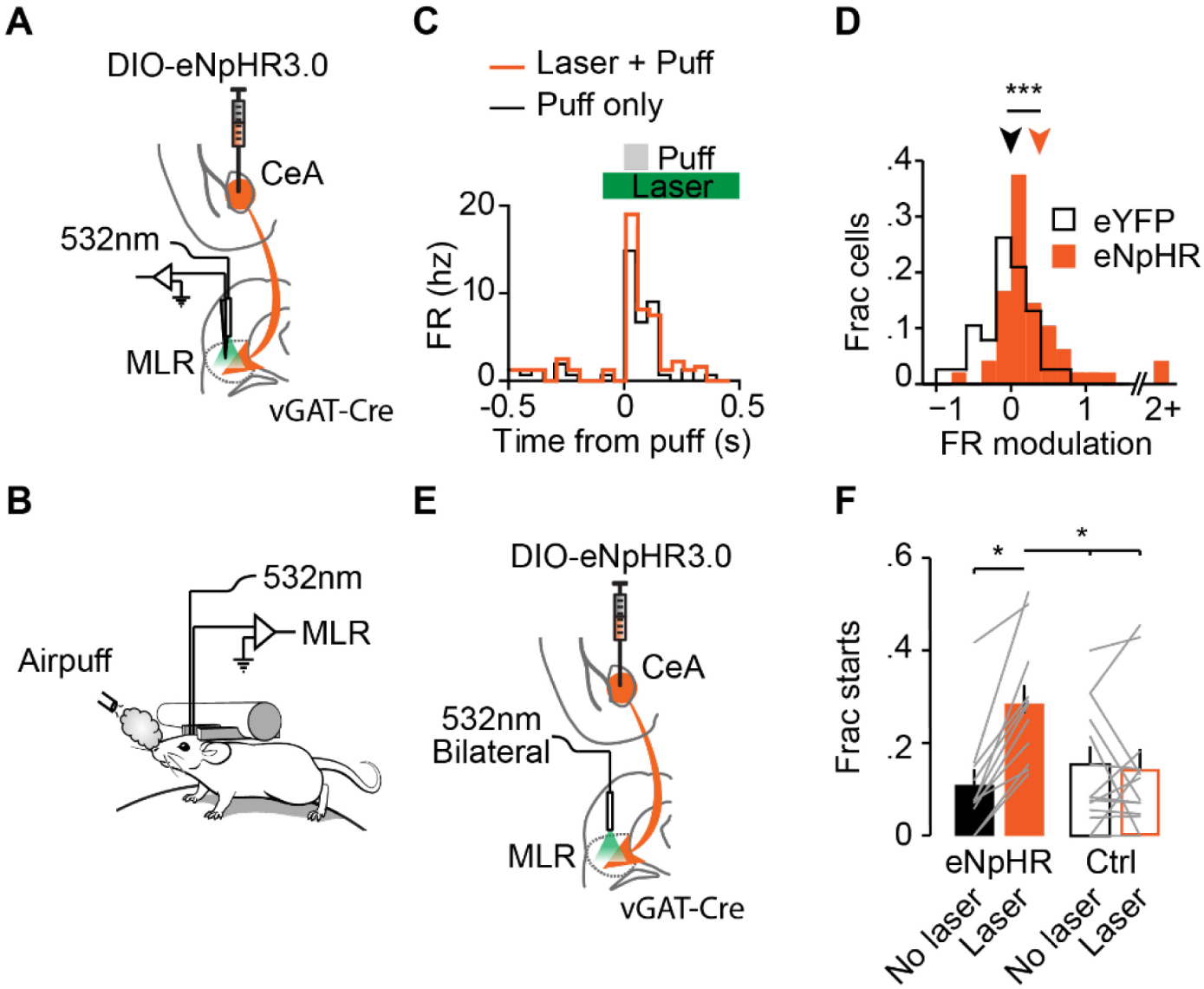
Inhibiting CeA terminals in the MLR increases firing and locomotion in response to an aversive stimulus. (A) Schematic of strategy for inhibiting CeA terminals while recording neurons in the MLR. (B) Experimental setup. (C) Example MLR neuron responding to airpuff with (orange) and without (black) laser. Green bar, laser duration. (D) Histogram of FR modulation during airpuff with and without laser for mice injected with eNphR3.0 (orange) or just eYFP (grey) (eNphR, n = 43 neurons from 5 mice; eYFP, n = 39 neurons from 4 mice; ***p < .001, rank-sum). (E) Schematic of strategy for inhibiting CeA terminals while recording neurons in the MLR. (F) Fraction of starts with (orange and orange outline) and without (black and black outline) laser (eNphR, n = 11 mice; eYFP, n = 14 mice; *p < .05, Two-way ANOVA χ^2^_3,61_ = 3.98, with Dunn-Sidak post test).

As bilateral inhibition of the MLR is required to modulate locomotion (Roseberry et al., 2016), in a separate set of mice we performed bilateral inhibition of CeA terminals in MLR while monitoring locomotion of head-fixed mice on a circular treadmill (Figure 4B & 4E). Optogenetic suppression of CeA projections to MLR significantly increased the probability that mice would start running within 1 second following an aversive airpuff, while eYFP controls showed no increase (Figure 4E-F). Together these results suggest that the CeA inhibits locomotion during aversive stimuli through regulation of MLR neurons.

### The CeA-MLR Pathway Inhibits Escape During Conditioned Avoidance

CeA projection neurons are active during conditioned avoidance and activating these neurons stops avoidance responses (Yu et al., 2016). Therefore, we wondered if the CeA-MLR pathway could be inhibiting conditioned avoidance responses. To investigate this, we performed three separate manipulations using the same task conditions to test: (1) if increased CeA-MLR activity could stop avoidance responses, (2) if MLR glutamatergic activity is necessary for avoidance responses, and (3) if inhibiting CeA-MLR terminals would increase avoidance responses or speed during the task. For all manipulations, virus was injected in either the CeA or MLR and fiber optics placed over the MLR (Figure 5D & 5G and Supplemental Figure 3A-E). After a 5 week recovery and expression period, mice were trained on a conditioned avoidance task (Yu et al., 2016) (Figure 5A - C). In the task, a head-fixed mouse was placed on top of a running wheel. Every 20 to 40 seconds (uniformly distributed), if the mouse was stationary, a white noise signalled that the mouse had to locomote above 5 cm/s in the next 1.5 seconds to avoid an airpuff otherwise lasting 5 seconds or until the mouse ran (Figure 5A-B). If the mouse ran during the white noise, no airpuff was delivered and the trial was considered successful. Mice increased the number of successful avoidances over multiple days (Figure 5C). After mice reached 50% correct trials for a day (Figure 5C), constant green laser for inhibition with eNpHR3.0 or 40 Hz blue laser for activation with ChR2 was delivered on 25% of trials for 4.5 seconds (Figure 5A).

**Figure 5.**
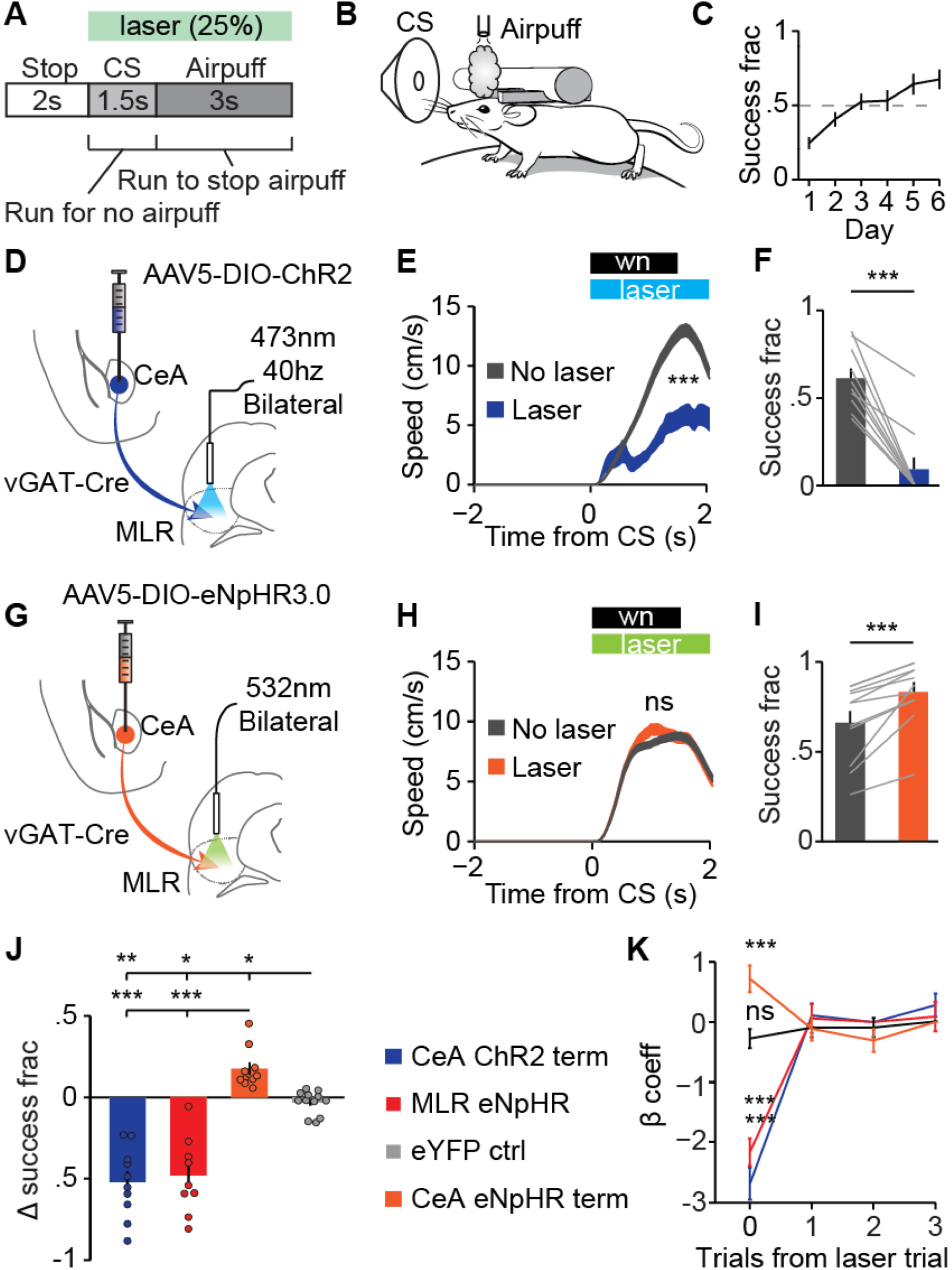
Manipulating CeA terminals and MLR^glut^ activity bi-directionally affects active avoidance performance. (A) Schematic of task with laser duration. (B) Schematic of CS and airpuff delivery. (C) Training time course. Laser trials begun after mice reached 50% success criteria (n = 46 mice). (D, G) Schematic for activating or inhibiting CeA terminals in the MLR, respectively. (E, H) Speed traces for trials with (colors) and without (grey) laser for CeA terminal excitation (E, n = 10 mice) or inhibition (H, n = 11 mice). Shaded traces, SEM (ns not significant, *** p <.001, rank-sum of average velocity with and without laser). (F, I) Fraction of successful trials with (colors) and without (grey) laser for CeA terminal excitation or inhibition (*** p < .001, rank-sum). (J) Summary of change in successful trials for all experimental conditions. (eYFP control n = 14 mice; *** p < .001, ** p < .01, * p < .05, Kruskal-Wallis one-way ANOVA χ^2^_3,45_ = 38.13, p < 10^−8^ with Dunn-Sidak post test). (K) Generalized linear model β coefficients for trial outcome after a stimulation trial (ns not significant, *** p < .001, χ^2^ test). Error bars, SEM.

We first asked if increased activity at CeA-MLR synapses could inhibit successful escapes. vGAT-Cre mice were injected bilaterally with DIO-ChR2 in the CeA and fiber optics were implanted over the MLR (Figure 5D and Supplemental Figure 3A). During laser trials, the percentage of successful escapes was significantly decreased, with many mice unable to perform a single successful trial (Figure 5F & 5J). When mice did escape, their speed was slower (Figure 5E). This demonstrates that the CeA-MLR connection can indeed inhibit successful escapes.

As the CeA-MLR pathway is predicted to control locomotion through suppression of MLR glutamatergic neurons, we next tested if activity in MLR glutamatergic neurons was required for successful escape. vGLUT2-Cre mice were bilaterally injected with DIO-eNpHR3.0 in the MLR and fiber optics were implanted over the injection site (Supplemental Figure 3B & 3E). Similar to terminal excitation, mice were significantly slower on successful trials in which the laser was on (Supplemental Figure 3F) and overall less likely to perform a successful escape (Figure 5J and Supplemental Figure 3G). This suggests that MLR glutamatergic activity is required for successful avoidance responses.

Finally, to test whether the CeA-MLR connection was preventing escape we inhibited CeA terminals in the MLR during the conditioned avoidance task. vGAT-Cre mice were injected in the CeA with DIO-eNpHR3.0, and optical fibers were implanted over the MLR (Figure 5G and Supplemental Figure 3C). While there was no significant effect on speed or reaction time during successful trials (Figure 5H), mice were on average 17% more likely to escape during a laser trial than a no laser trial (Figure 5I and 5J). Interestingly, none of these manipulations had a significant effect on subsequent trials (Figure 5K) indicating that there was no learning induced. This indicates that the CeA-MLR connection is inhibiting escape responses to learned aversive stimuli, similar to our findings obtained with an innately-aversive noxious stimulus (Figure 4).

## Discussion

In the present study, we delineate a circuit mechanism for suppression of defensive locomotion in response to aversive stimuli. GABAergic CeA neurons suppress locomotion via a monosynaptic connection with glutamatergic neurons in the MLR.

### A mechanism for specialized locomotor suppression in response to aversive stimuli

Neurons within the MLR are spontaneously active *ex vivo* (Granata and Kitai, 1991; Kang and Kitai, 1990). This activity is inhibited tonically by local GABAergic neurons (Roseberry et al., 2016), as well as inhibitory afferents from the Substantia Nigra pars reticulata (SNr) (Granata and Kitai, 1991; Grillner et al., 2008; Kang and Kitai, 1990). Consequently, disinhibition of MLR by suppression of these tonically active inhibitory inputs is critical for locomotion (Garcia-Rill, 1986; Garcia-Rill et al., 1985; Hikosaka et al., 2000; Nandi et al., 2002). Surprisingly, we found that the CeA-MLR connection does not contribute to this tonic inhibition and does not regulate spontaneous locomotion. Instead, GABAergic input from the CeA specifically shapes locomotor responses to aversive contexts.

In addition to potential disinhibitory mechanisms, the burst of activity in the MLR in response to an aversive airpuff stimulus (Figure 1) likely arises from a phasic excitatory input. Candidate sources for this excitatory input include the ventral thalamus (Jordan, 1998; Menard and Grillner, 2008) and zona incerta (Menard and Grillner, 2008). Both regions have been proposed to constitute the diencephalic locomotor region (DLR) where electrical stimulation can also drive locomotion (Menard and Grillner, 2008; Sinnamon, 1993). Both regions are also innervated by the basal ganglia (BG) (Menard and Grillner, 2008) which is implicated in learned motor behaviors and is required for successful active avoidance (Hormigo et al., 2016; Vecsei and Beal, 1991). Together, the CeA and DLR could send competing inhibitory and excitatory inputs, respectively, to the MLR (Jordan, 1998). The timing and dynamics of this competition and the exact site of the upstream excitatory input are intriguing future questions.

### Anatomy of the CeA-MLR projection

It is interesting to note that the MLR has recently been divided into two parts: a ventral portion mainly comprised of the dorsal PPTg, and a dorsal portion comprised of the Cuneiform nucleus (Caggiano et al., 2018). The cuneiform neurons were demonstrated to be necessary for controlling faster escape-type running speeds while the PPTg neurons supported slower exploratory locomotion. Our anatomical tracing and physiological results found that the CeA innervates both structures suggesting that the CeA signal could be a global stop signal for all types of locomotion. Indeed, as MLR neurons are spontaneously active (Kang at al., 1990), suppressing both the dorsal and ventral MLR would be necessary for freezing and bracing for flight – both crucial defensive behaviors.

### Anatomical and cellular diversity within the CeA

Our results indicate that the MLR receives monosynaptic inhibition from both the centrolateral (CeL) and centromedial (CeM) subregions of the central amygdala (CeA). The absence of an apparent distinction between CeL and CeM in this output projection to MLR may appear surprising in light of evidence for a functional opposition between CeL and CeM, in which CeL neurons inhibit projection neurons in the neighboring CeM (Ciocchi et al., 2010; Fadok et al., 2017; Huber et al., 2005; Li et al., 2013; Viviani et al., 2011). However, tracing studies from the PAG suggest that the distinction between CeL and CeM may be less clear-cut in mouse (Tovote et al., 2016) than in rat (Viviani et al., 2011). In addition, the CeL houses a heterogeneous population of neuronal subtypes, at least some of which are functionally aligned with CeM activity (Cui et al., 2017; Fadok et al., 2017; Li et al., 2013; Yu et al., 2016). It is interesting to note that somatostatin neurons in the CeA respond mainly to conditioned stimuli and can inhibit conditioned avoidance responses (Cui et al., 2017; Yu et al., 2016), but do not respond to airpuff (Cui et al., 2017). However, CeA PKCδ-expressing neurons do respond to airpuff (Cui et al., 2017). It is possible that both somatostatin and PKCδ neurons project to the MLR and inhibit MLR neurons in different contexts – somatostatin for conditioned responses and PKCδ for unconditioned responses. Future studies will be required to address this interesting question.

### Behavioral relevance of the CeA to MLR projection

The freezing response to fear-inducing or aversive stimuli is a highly stereotyped and rigidly defined behavior. In addition to facilitating freezing by suppressing potentially competing motor plans, the CeA-MLR circuit may also play a role in a less stereotyped “deer in the headlights” response, or failure to take action when a surprising or known dangerous stimulus is presented. While it may seem detrimental for an animal’s survival to inhibit locomotion upon detection of a dangerous stimulus, a pause before action could allow for a more complex motor plan to develop (Gladwin et al., 2016; Roelofs, 2017). Simple reflexive escape responses could result in running toward the dangerous stimulus or into stationary objects. A pause could allow assessment of where the danger is, where the nearest safe spot is, what barriers might lie in the way, and whether or not freezing or running is the best option given the proximity to danger. Indeed threat proximity, stimulus size, and shape all change rodent behavior from freezing, when a threat is further away, to escape, if the threat is near or looming (De Franceschi et al., 2016). The CeA, which is active during both learned and innately aversive stimuli (Cui et la., 2017; Yu et al., 2016), could provide this pause signal, but may also induce muscle rigidity to prime the animal for escape through a CeA-PAG-Pontine pathway recently demonstrated by Tovote et al., 2016. Through a series of excitatory and disinhibitory connections, the CeA-PAG-Pontine pathway excites motor neurons in the spinal cord. In coordination with the projection to the MLR, the overall effect would be to inhibit locomotion while maintaining muscle tone for faster initiation of locomotion, should escape be required. It is still unclear if all MLR-projecting CeA neurons are also PAG-projecting neurons. Either way, many CeA projection neurons appear to be activated by a conditioned stimulus (Ciocchi et al., 2010; Yu et al., 2016) and could thus coordinate a pause and escape response as single or separate populations.

The results presented here expand the circuit substrates for defensive behaviors and provide critical insights into why both animals and humans can fail to take action when placed in a dangerous or high stress situation. As the number of people affected by fear-based disorders grows (Luthi and Luscher, 2014), a mechanistic understanding of how the underlying circuitry functions will prove crucial to formulating treatments and implementing preventions.

**Supplemental Figure 1.**
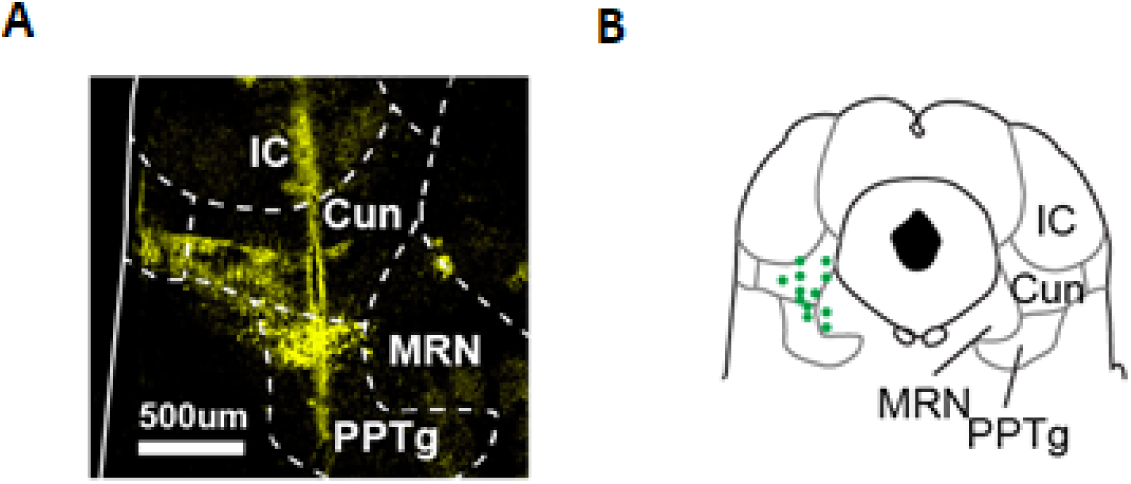
MLR glutamatergic identified recording locations. Related to Figure 1. (A) Example electrode track in MLR. (B) Recording locations for MLR glutamatergic identified neurons.

**Supplemental Figure 2.**
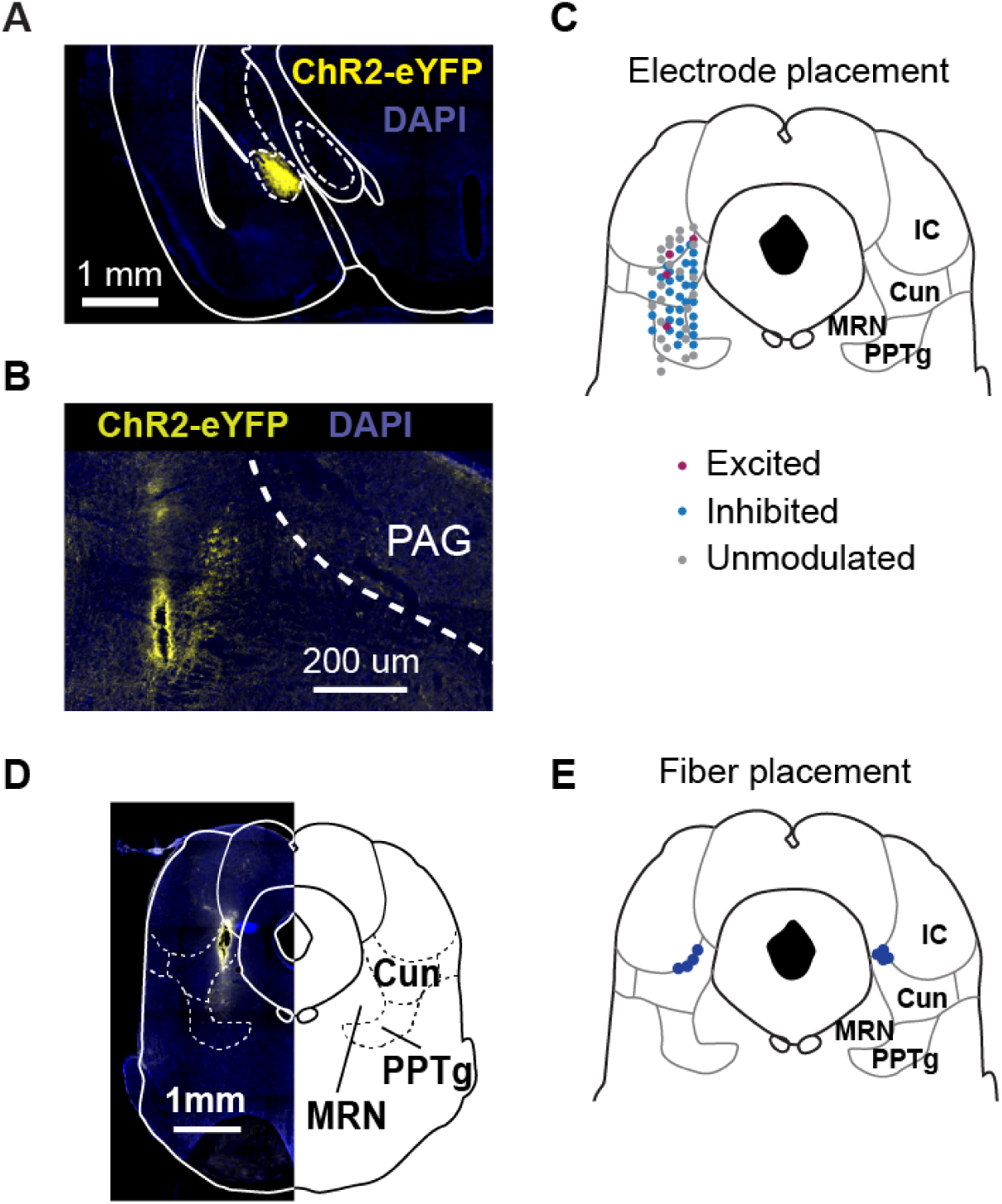
Viral injection, recording and fiber placements. Related to Figure 2 and 3. (A) Example injection in the CeA. (B) Example recording track in the MLR. (C) Schematic of recording locations from each drive with cell response to CeA terminal stimulation listed. (D) Example fiber track locations. (E) Schematic of all fiber track locations.

**Supplemental Figure 3.**
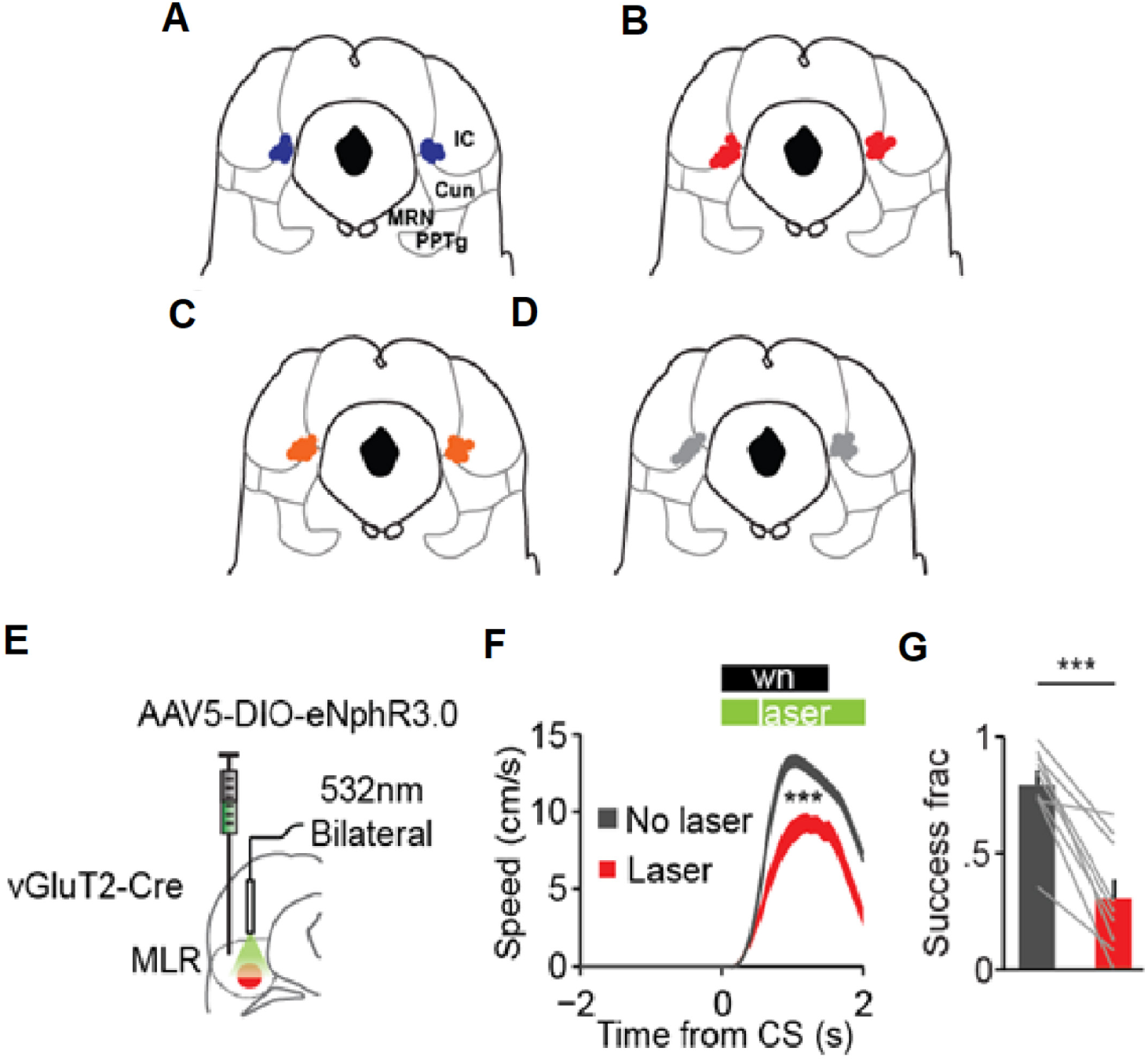
Active avoidance fiber placement and inhibition of MLR glutamatergic neurons during active avoidance. Related to Figure 5. (A - D) Placement of optical fiber tips in the MLR for (A) CeA terminal excitation using ChR2. MLR glutamatergic inhibition using eNpHR3.0, (C) CeA terminal inhibition using eNpHR3.0, (D) eYFP controls. (E) Schematic for inhibiting MLR glutamatergic neurons using eNpHR3.0. (F) Results of inhibiting MLR glutamatergic neurons on 25% of trials (”Laser”) vs no inhibition (”No laser”) (n = 9 mice, *** p < .001 sign-rank for average velocity during CS). (G) Change in fraction of successful trials during laser inhibition (*** p < .001, sign-rank). Error bars, SEM.

## Methods

### Key Resources

| REAGENT or RESOURCE | SOURCE | IDENTIFIER |
| --- | --- | --- |
| Bacterial and Virus Strains |  |  |
| CAV2-Cre | IGMM | n/a |
| AAV5-EF1α-ChR2-eYFP | U Penn Vector Core | n/a |
| AAV5-EF1α-eNpHR3.0-eYFP | U Penn Vector Core | n/a |
| AAV5-EF1α-eArch3.0-eYFP | U Penn Vector Core | n/a |
| AAV5-EF1α-eYFP | U Penn Vector Core | n/a |
| AAV1-Syn-Flex-GCaMP6m-WPRE-SV40 | U Penn Vector Core | n/a |
| Experimental Models: Organisms |  |  |
| Mouse: C57BL/6 | Jackson Laboratories | 000664 |
| Mouse: vGLUT2-ires-Cre | Jackson Laboratories | 016963 |
| Mouse: vGAT-ires-Cre | Jackson Laboratories | 016962 |
| Mouse: Allen Institute line Ai14(RCL-tdT)-D | Jackson Laboratories | 007914 |
| Software and Algorithms |  |  |
| MATLAB | Mathworks | Mathworks.com |
| NeuroExplorer V4.133 | Neuroexplorer | Neuroexplorer.com |
| Plexon Offline Sorter V2.1 | Plexon Inc | Plexon.com |
| Fiji | ImageJ | Fiji.sc |

### Animals and stereotactic surgery

vGLUT2-ires-Cre (Jackson Stock# 016963), vGAT-ires-Cre (Jackson Stock# 016962), Allen Institute line Ai14(RCL-tdT)-D (Jackson Stock# 007914) and C57BL/6 (Wild-type, Jackson Stock# 000664) mice were used for experiments where noted. Mice were 50 to 55 days old when entering surgery. All procedures were in accordance with protocols approved by the UCSF Institutional Animal Care and Use Committee. Mice were maintained on a 12/12 light/dark cycle and fed *ad libitum*. Experiments were carried out during the dark cycle. All surgeries were carried out in aseptic conditions while mice were anaesthetized with isoflurane (5% for induction, 0.5-1.5% afterward) in a manual stereotactic frame (Kopf). Buprenorphine HCl (0.1 mg kg^-1^, intraperitoneal injection) and ketoprofen (5 mg kg^-1^, subcutaneous injection) were used for postoperative analgesia.

### Viral injections

For cell-type-specific viral infection of CeA neurons, we injected 150 to 200 nL of adeno-associated virus serotype 5 (AAV5) carrying channelrhodopsin 2.0 (ChR2), enhanced halorhodopsin 3.0 (eNpHR3.0) or enhanced archaerhodopsin 3.0 (eArch3.0) fused to enhanced yellow fluorescent protein (eYFP) in a double-floxed inverted open reading frame (DIO) under the control of the EF1α promoter. AAV5-EF1α-DIO-eYFP was used as a fluorophore-only control virus. CeA coordinates were −1.2 mm anteroposterior (AP) from bregma, ± 1.2 mm mediolateral (ML) and −4.8 mm dorsoventral (DV) from skull surface. For inhibition or activation of glutamatergic neurons in the MLR, we injected 300 nL of AAV5-EF1α-eNpHR3.0-eYFP or AAV5-EF1α-ChR2-eYFP, respectively, in vGLUT2-Cre mice. MLR injections were made bilaterally at + 0.2 mm from the Lambda suture, ± 1.2 mm ML and −3.4 mm DV measured from the skull surface. For labelling MLR-projecting CeA neurons, 200 nL of CAV2-Cre virus (Institut de Génétique Moléculaire de Montpellier) was injected unilaterally into MLR coordinates of Ai14 mice. For photometry experiments 200 nL of CAV2-Cre was injected unilaterally into the MLR of C57BL6 mice followed by 200 nL of AAV1-Syn-Flex-GCaMP6m-WPRE-SV40 into the CeA. Injections were made using a 5 µL NanoFil with 35ga Needles (WPI) mounted to a microsyringe pump (UMP3; WPI) and controller (Micro4; WPI). Injection speed was 50 nL min^-1^ and the injection needle was raised 7-10 minutes after completion.

### Preparation for Head Fixation

For all experiments, mice were implanted with a custom stainless steel headbar for head fixation. For acute recordings, this took place 3 to 4 weeks after viral injection. For photometry recordings or optogenetic manipulation experiments, this took place during the viral injection surgery. The scalp was removed and the skull scraped clean and dry using a scalpel. Cyanoacrylate glue (Vetbond, 3M) or mixed Metabond C&B quickbase powder and catalyst (Parkell) was lightly dabbed on the skull and a base of dental acrylic was applied in a circle around Lambda to provide the skull/headbar interface (Ortho Jet; Lang Dental). For acute recordings a headbar with a 6 mm hole was placed around the future recording area, further cemented, and the hole then filled with silicone elastomer (Kwik-Cast, WPI). After 7 to 10 days of recovery, a 0.5 mm burr hole was drilled above the recording region (MLR or CeA). The headbar was then again filled with silicone elastomer and the animal allowed to recover for 3 to 4 hours after which it was placed on the running wheel for recording experiments. For photometry recordings a 500 um hole was drilled above the recording site at the AP and ML CeA coordinates. A 400 um polished fiber glued into a ferrule sleeve (Precision Fiber MM-FER2003SS-4500) was then lowered to 4.5 mm DV from brain surface over 30 minutes. For optogenetic experiments, 200-µm-diameter optical fibers (Thorlabs #FT200UMT) were implanted bilaterally over the MLR at a DV of 2.9 mm after headbar mounting by drilling a 0.5 mm hole in the skull, lowering the fiber over 1 minute and gluing into place with Vetbond, followed by Metabond and dental acrylic.

### Untrained Behavior and Recording Parameters

For spontaneous locomotion behavior, mice were habituated to the wheel for 2 to 3 days prior to the experiment. Speed was tracked via a quadrature encoder and custom MATLAB software. For optical stimulation experiments, blue or green light was passed through a 200 µm fiber (described below) using a 473 nm or 532 nm laser (OEM part # OEM-S-473 or OEM-S-532) coupled to an optical multimode fiber (200 µm, 0.39 NA FC/PC, Thorlabs part # M83L01). For recording experiments, the light was passed through a 200 µm fiber attached to a recording probe (described below). Blue light for terminal excitation was delivered at 5 to 7 mW at the fiber tip at 40 Hz in 25 ms pulses triggered via TTL pulses produced from a wave generator (TFNMA part# TGP110). Green light for terminal inhibition was delivered continuously at 7 to 8 mW at the fiber tip and shuttered (Uniblitz VCM-D1) to maintain power stability. The shutter and laser were placed in a custom-built sound-proof box to block shutter clicks and triggered via the data acquisition box (Plexon).

For recording MLR responses to CeA terminal excitation, 10 ms light pulses were pulsed through a fiber attached to the electrode at 0.5 Hz. For terminal stimulation during running, custom closed-loop control software ensured the mouse was running for 5 seconds continuously prior to stimulation onset of the 5 second 40 Hz pulses. For CeA terminal inhibition, green light was delivered every 20 seconds regardless of running state.

To observe the effects on locomotion and firing during inhibition of CeA terminals in the MLR or while exciting glutamatergic neurons below a threshold that would elicit locomotion in response to airpuff, a solenoid (NResearch part# 161P011) was triggered for 100 ms using a PulseBlaster TTL generator (SpinCore) to release pressured air. Airpuff inter-trial-interval was 20 seconds fixed. Lasers were activated for 1 second starting at the onset on interleaved trials.

### Conditioned Avoidance Task

Mice were given one 30 minute habituation session prior to start of the task. For photometry experiments, mice were given one extra day for two separate conditions. In the first condition, the mouse was allowed to run freely for 40 minutes without distraction while signal was recorded, in order to establish a “spontaneous locomotor” condition. In the second condition run after the spontaneous condition, 100 CS (1.5 s white noise at 65 dB) were randomly delivered at 20 to 40 s intervals (ITI) without airpuff (US) while signal was recorded to establish “CS Naïve” response. The next day mice began the task. Mice undergoing optogenetic inhibition or excitation performed 200 trials each day, whereas photometry mice performed 100 to avoid phototoxicity and loss of GCaMP signal. Interestingly, there was no difference in learning rate between these groups despite trial difference. During the task, the CS was delivered when the mouse was stopped for 2 continuous seconds. If the mouse did not run during the 1.5 s WN, it would receive an airpuff directly to the eye. The airpuff would last for 3 s or when the mouse ran, whichever occurred first. However, both conditions were identified as failure trials. Custom MATLAB software was used to measure mouse speed and deliver TTL signals for the tone, laser, and airpuff from a NIDAQ-6009 data acquisition device (National Instruments). ITI was 20 to 40 s, evenly distributed. For stimulation or inhibition experiments, the day after a mouse completed 50% successful trials (2-6 days), laser stimulation trials were added for a single day. Laser was delivered from tone onset to 4.5 s regardless of trial outcome on 25% of trials. For CeA terminal inhibition or MLR glutamatergic neuron inhibition, 532 nm continuous light at 7-8 mW was delivered. For terminal excitation 473 nm light at 5-7 mW, 40 Hz with a 25 ms pulse, controlled by the wave generator, was delivered.

### In Vivo Electrophysiology

Extracellular spikes were recorded using NeuroNexus silicon probes (part no A1×16-10mm-100-177-A16). A 200-µm-diameter optical fiber was manually mounted on the probe, with the upper-most recording site approximately 100 µm below the tip of the fiber and 200 µm lateral from the probe surface. Once the probe had been lowered in the brain, a drop of agarose was used to stabilize it. Voltage signals were band-pass filtered between 150 and 8000 Hz. Each data stream was amplified, processed and digitally captured using commercial hardware and software (Plexon). Typically, only 1 unit was identifiable per recording depth. After each recording, the probe was driven 100 to 200 µm to find other cells. Neurons that appeared to stay on the same channel or appeared move up the probe sites as the probe was lowered were excluded.

For recording optogenetically-identified glutamatergic MLR neurons, a similar procedure was used as Roseberry et al., 2016. Once a stable recording was established, blue light was flashed for 10 ms at 2 to 10 mW through the fiber into the brain at 1-2 Hz for 200 to 300 repetitions. Laser power was adjusted to minimize the latency of activation while also minimizing optical artifact. Once a neuron was identified as light-sensitive, the airpuff session would proceed after which a post identification session would be carried out to ensure another unit had not moved into the recording space. Final clustering was performed post hoc.

### Analysis of Neural Data

Pre- and post-identification sessions were merged with the associated locomotor or stimulation session to sort single units on the same principle component and peak-to-trough amplitude spaces. Single units were sorted using Offline sorter V2.1 (Plexon). Recorded units in which more than 1% of interspike intervals were shorter than 2 ms were excluded from analysis. MANOVA (p < 0.01) and J3 (> 3.0) parameters in 3 dimensions were used to ensure cluster quality.

To determine excited, inhibited or unmodulated populations, the number of spikes recorded during each laser pulse (laser period) were compared to the number of spikes in the same amount of time prior to laser pulse onset (baseline period) using Wilcoxon rank sum. Firing rate modulation was determined by taking the average firing rate in the laser period and dividing it by the average firing rate in the baseline period. Latencies were determined by creating 1 ms bins in NeuroExplorer V4.133 and finding the first of 3 bins that were outside the 97% confidence interval.

For identifying MLR glutamatergic neurons, light responses were determined using 1 ms bins in NeuroExplorer V4.133. All other analysis of neural and behavioral data was carried out using custom MATLAB software (available upon request). A neuron was considered identified (ChR2-positive) if: (1) its firing rate cleared the 99% confidence interval (based on a 20 ms baseline period prior to stim) within 5 ms after the onset of the 10 ms pulse, (2) light-evoked spike waveforms (occurring within the 10 ms of the light pulse) were identical to the spontaneous waveform (R > 0.9), and (3) light-evoked responses displayed jitter relative to the light onset (indicating spikes were not related to a light artifact).

### Electrophysiology in acute slices

Mice were euthanized with a lethal dose of ketamine and xylazine followed by transcardial perfusion with 8 ml of ice cold artificial corticospinal fluid (aCSF) containing (in mM): glycerol (250), KCl (2.5), MgCl2 (2), CaCl2 (2), NaH2PO4 (1.2), HEPES (10), NaHCO3 (21) and glucose (5). Coronal slices (250μM) containing the MLR were then prepared with a vibratome (Leica) in the same solution, before incubation in 33°C aCSF containing (in mM): NaCl (125), NaHCO3 (26), NaH2PO4 (1.25), KCl (2.5), MgCl2 (1), CaCl2 (2), glucose (12.5), continuously bubbled with 95/5% O_2_/CO_2_. After 30 minutes of recovery, slices were either kept at room temperature or transferred to a recording chamber superfused with recording aCSF (2.5 ml/min) at 33°C. Voltage-clamp recordings were obtained using an internal solution containing (in mM): CsMeSO3 (15), CsCl (120), NaCl (8), EGTA (0.5), HEPES (10), Mg-ATP (2), Na-GTP (0.3), TEA-Cl (10), QX-314 (5). Synaptic release was evoked by flashing 470nm filtered LED (Prizmatix) blue light through the objective (1-10 msec pulse, 1mW/cm^2^). For connectivity from the CeA, cells were considered not connected if no IPSC was detectable with 1.5 mW of illumination. IPSCs were measured at 20s intervals, while holding the membrane potential at −60 mV. The MLR was identified as the region lateral to the decussation of the superior cerebellar peduncle. Picrotoxin waspurchased from Tocris, prepared at stock concentration in H_2_0, then diluted in aCSF for bath application. Data was acquired with Igor software.

### Histology

Animals were euthanized with a lethal dose of ketamine and xylazine (400 mg ketamine plus 20 mg xylazine per kilogram of body weight, i.p.) and transcardially perfused with PBS, followed by 4% paraformaldehyde (PFA). Following perfusion, brains were transferred into 4% PFA for 16 –24 h and then moved to a 30% sucrose solution in PBS for 2–3 d (all at 4 deg C). Brains were then frozen and cut into either 100 or 30 µm coronal sections with a sliding microtome (Leica Microsystems, model SM2000R) equipped with a freezing stage (Physitemp) and mounted on slides. For confirmation of fiber tracks and viral injection, slices were quickly imaged and photographed under an Olympus BX51WI microscope. Pertinent slices were then kept for mounting on slides.

Slides were blocked for 1 hour in 10% Normal Donkey Serum (NDS) in 0.5% PBST then incubated overnight in primary antibody (1:500), 3% NDS in 0.5% PBST. The following day, they were washed 3 times for 10 minutes each in 0.5% PBST and incubated for 1 hour in secondary antibody (1:1000), 3% NDS in 0.5% PBST and 1:2000 DAPI. After this, slides were washed for 10 minutes in 0.5% PBST and 2 more 10 min periods with 1:1 PBS. Slides were then washed with 0.05% lithium carbonate and alcohol, rinsed with diH2O, and coverslipped with Cytoseal 60.

Figure images were acquired using a 6D high throughput microscope (Nikon, USA) or SP5 confocal (Leica, USA), globally gamma-adjusted to reduce background, and pseudocolored using freely available Fiji software. To determine recording sites along the dorsal-ventral axis, the lowest depth of the electrode was noted both post hoc and during the experiment. The difference between these two values was then subtracted from the noted depth during a given recording. For recordings, the recording site distance from the electrode tip was further subtracted to give a more exact position. The electrode track itself was used to locate the ML and AP position. Neurons determined to be outside of the MRN, Cun or PPTg were excluded.

### Statistics

All statistics are noted in figure legends. Non-parametric tests were used in all cases except 2-Way ANOVAs in which case data was tested for normality using the Lillie Test and an α of 0.05.

Data and code are available on request.

## Acknowledgements

We thank the Institut de Génétique Moléculaire de Montpellier for providing CAV2-Cre virus. We thank the Kreitzer lab for assisting with experimental setups and thoughtful discussion. This work was funded by NIH R01 NS064984 to A.C.K., F31 NS092253 to T.K.R., RR018928 to the Gladstone Institutes, and a grant from the Swiss National Science Foundation to A.L.L.

## Author Contributions

T.K.R. and A.C.K. conceptualized the project, designed all experiments, analyzed data and wrote the manuscript. A.L.L performed and analyzed slice electrophysiology experiments, and B.D.M. performed and analyzed some experiments.

## Declaration of Interests

The authors declare no competing interests.

